# Establishment of primary renal lymphoma model and the clinical relevance

**DOI:** 10.1101/2023.01.03.522576

**Authors:** Xiaoxi Li, Minyao Deng, Chenxiao Zhang, Lingli Luo, Hui Qian

**Affiliations:** Department of Laboratory Medicine, School of Medicine, Jiangsu University, Zhenjiang, Jiangsu, China

**Keywords:** Primary renal lymphoma, Extranodal lymphoma, Extranodal dissemination, MA-K, MCD subtype, LymphGen, Aggressive B-cell lymphoma, Translation pathway

## Abstract

Extranodal dissemination was an important feature of aggressive B-cell lymphoma. Due to the lack of available animal model, the causes of extranodal dissemination of lymphoma were largely unknown. Here, we identified a novel cell line, named MA-K, which originated from the Eμ-Myc;Cdkn2a^-/-^ cell line, named MA-LN in this study. Compared to MA-LN, MA-K tended to disseminate in the kidney rather than lymph nodes in lymphoma transplantation model, resembling human primary renal lymphoma. The transcriptome analysis revealed that MA-K had undergone the transcriptional evolution during the culture. Specialized transcriptional pattern analysis, we proposed in this study, identified that FOXO1-BTG1-MYD88 pattern was formed in MA-K. Further analysis found that translation pathway was the most enriched pathway in specially expressed genes (SEGs) in MA-K. Among the SEGs, 3 up-regulated genes, RPLP2, RPS16 and MRPS16, and 5 down-regulated genes, SSPN, CD52, ANKRD37, CCDC82 and VPREB3, in MA-K were identified as promising biomarkers to predict the clinical outcomes of human DLBCL. Moreover, the joint expression of the 5-gene signature could effectively predict clinical outcomes of human DLBCL in triple groups. These findings suggested that the MA-K cell line had strong clinical relevance with human aggressive B-cell lymphoma. Moreover, the MA-K primary renal lymphoma model, as a novel syngenetic mouse model, would be greatly useful for both the basic research on lymphoma dissemination and preclinical efficacy evaluation of chemotherapy and immunotherapy.

## Introduction

B-cell lymphoma was a B-lymphoid hyperplasia disease with high heterogeneity. While most lymphomas primarily presented in lymph nodes, extranodal dissemination of lymphoma was a common clinical feature observed in most subtypes of Non-Hodgkin’s B-cell lymphoma (B-NHL), including diffused large B-cell lymphoma (DLBCL)(1,2), Burkitt’s lymphoma (BL)(3,4), high-grade B-cell lymphoma (HGBL)(5,6) and others. The disseminated organs included the central nervous system (CNS)(7), skin(8), uterine(9). Primary renal lymphoma (PRL) was a rare malignant lymphoma and most of PRL cases were DLBCL(10). Patients with extranodal lymphoma, such as CNS lymphoma, often had poor clinical outcome. Classification of DLBCL based on transcriptional profile(11) and genetic variation(2) had link the extranodal lymphoma to activated B-cell-like (ABC) subtype and MCD (including MYD88^L265P^ and CD79B mutations) subtype. However, due to the lack of available animal models, the genetic and non-genetic factors of extranodal lymphoma were still unclear.

Eμ-Myc transgenic mouse was a well-established spontaneous B-cell lymphoma mouse model(12) resembling the translocation of oncogenic Myc to the enhancer of immunoglobulin heavy (IgH) μ gene in human BL. Unlike human BL and DLBCL originated from mature B stage, the later stage of B-cell differentiation, Eμ-Myc lymphoma mainly originated from pro-B and pre-B stage. Hence, Eμ-Myc transgenic mouse was not enough to resemble the aggressive phenotype of human B-cell lymphoma, such as extranodal dissemination. In the combined Eμ-Myc transgenic mouse and genetically engineered modified mouse (GEMM) models, the knockout of tumor suppressor genes (TSGs), such as p53 and Arf (13,14), could significantly accelerate lymphomagenesis and shorten survival time. Transcriptome analysis on a large cohort of Eμ-Myc transgenic mice(15) revealed that the onset of Eμ-Myc lymphoma dramatically varied and BL-like and DLBCL-like transcriptional characteristics were identified in early-onset and late-onset lymphoma respectively. In addition, genomic analysis (16) identified that disruptive mutations in Bcor contributed to spontaneously lymphomagenesis of Eμ-Myc transgenic mouse. Given that lymph nodes were still the major disseminated sites in most Eμ-Myc based mouse models, understanding the mechanism of extranodal dissemination was still difficult and challenging.

Due to easy to establish syngeneic lymphoma transplantation model, a kind of the GEM-derived allograft (GDA) model(17), Eμ-Myc;Cdkn2a^-/-^ cell line, usually giving rise to lymphoma in lymph nodes, was widely used for in vivo efficacy evaluation(18,19). In this study, we reported a Eμ-Myc;Cdkn2a^-/-^ derived cell line that could give rise to extranodal lymphoma, specifically to kidney, in GDA model. To distinguish with parental Eμ-Myc;Cdkn2a^-/-^ cell line, we named the kidney-disseminated cell line as MA-K cell line, in which the M referred to Myc and the A referred to Arf. Since the strong clinical relevance of extranodal dissemination and aggressive B-cell lymphoma, we further analyzed the transcriptome profile of MA-K and explored prognostic biomarkers of human DLBCL inspired by the transcriptome of MA-K. Translation pathway was the most enriched pathway in SEGs in MA-K. 8 SEGs in MA-K, RPLP2, RPS16, MRPS16, SSPN, CD52, VPREB3, CCDC82, ANKRD37, were identified as promising prognostic biomarkers of human DLBCL. Together, we reported a novel MA-K cell line that tended to kidney dissemination in GDA model had strong clinical relevance with aggressive DLBCL. The MA-K GDA model should be a useful model to explore the genetic and non-genetic mechanism of primary renal lymphoma and preclinical efficacy evaluation for both chemotherapy and immunotherapy.

## Materials and methods

### Cell lines

The *Eμ-Myc;Cdkn2a^-/-^* cell line, also named MA-LN in this study, was a kind gift from Prof. Michael Hemann at MIT in 2011 and preserved in the lab of Prof. Hai Jiang at CEMCS, CAS. The MA-K cell line was established from the *Eμ-Myc;Cdkn2a^-/-^* cell line in our lab. The *Eμ-Myc;Cdkn2a^-/-^* cell line and the MA-K cell line were cultured in 45%DMEM, 45%IMDM, 10% Fetal Bovine Serum (Biosera, FB-1058), supplemented with 100 U/ml penicillin and streptomycin, 25μM β-mercaptoethanol.

### Lymphoma transplantation model

All mice were housed in a specific pathogen-free environment at Laboratory Animal Research Center in Jiangsu University and treated in strict accordance with protocols, which were approved by the Animal Care and Use Committee of Laboratory Animal Research Center, Jiangsu University.

6-week C57BL/6JGpt female mice were purchased from the GemPharmatech (Nanjing, CN). 10^6^ MA-LN cells or MA-K cells in 200μl of DPBS were injected into C57BL/6 female recipient mice via tial vein. Recipient mice transplanted with MA-LN cell line usually grown palpable mass at axillary lymph nodes at 4th week post of transplantation. Instead, typical symptoms, including hunch back, dull hair, movement retardation and abdominal bulge, usually were observed in MA-K recipient mice at 4th week post of transplantation. Recipient mice transplanted with MA-LN or MA-K were monitored for any one symptom appeared and were sacrificed for evaluating lymphoma dissemination.

### H&E staining and imaging

For histological H&E staining, lymphomas were fixed in 10% formalin overnight and subsequently transferred into 70% ethanol, embedded in paraffin according to standard protocols. Sections (8 μm) were stained with H&E and Images from the whole slide were acquired by Pannoramic MIDI (3DHISTECH) and analyzed by CaseViewer software (3DHISTECH).

### RNA extraction and RNA sequencing

Two replicates of MA-LN and MA-K, collected at different time, were applied to RNA sequencing. Total RNA was extracted using Trizol reagent kit (15596018, Invitrogen) according to the manufacturer’s protocol. RNA library construction and sequencing were performed by Gene Denovo Biotechnology Co. (Guangzhou, China). The enriched mRNA by Oligo(dT) beads was fragmented into short fragments using fragmentation buffer and reversly transcribed into cDNA by using NEBNext Ultra RNA Library Prep Kit for Illumina (#7530, New England Biolabs). The purified double-stranded cDNA fragments were end repaired, A base added, and ligated to Illumina sequencing adapters. The ligation reaction was purified with the AMPure XP Beads (1.0X). Ligated fragments were subjected to size selection by agarose gel electrophoresis and polymerase chain reaction (PCR) amplified. The resulting cDNA library was sequenced using Illumina Novaseq6000.

### Specially expressed genes (SEGs) analysis

For each transcription region, a FPKM (fragment per kilobase of transcript per million mapped reads) value was calculated to quantify its expression abundance and variations, using RSEM software.

To filter SEGs with biological significance, we divided the gene expression level into 3 levels as FPKM>=10, FPKM>=1, FPKM<1. Inactive genes or basal expressed genes were defined as FPKM<1. Active genes were defined as FPKM>=1. SEGs were filtered as follows. Inactive genes (FPKM<1) in both groups were directly excluded for SEGs analysis. For active genes (FPKM>=1) in both groups, genes with Log_2_(FC)<=-2 and Log_2_(FC)>=2 were filtered as SEGs. In the case of inactive genes (FPKM<1) in one of groups, active genes (FPKM>=10) in another group were directly listed into SEGs. All SEGs were listed in Data Sheet 1.

Pathway enrichment analysis and protein-protein interaction enrichment of SEGs were performed with Metascape (https://metascape.org). Protein-protein interaction enrichment analysis had been carried out with the following databases STRING, BioGrid. Only physical interactions in STRING (physical score > 0.132) and BioGrid were used.

### Survival analysis

Survival analysis was performed with the online tool SurvExpress (20). A human DLBCL dataset (Lenz Staudt Lymphoma GSE10846(21), n = 420) was chosen to survival analysis. The prognostic index (PI) was calculated by the expression value and the Cox model to generate the risk groups. The optimization algorithm was applied in risk grouping. The SurvExpress program was performed according to the tutorial.

### Statistical analysis

Depending on the type of experiment, log-rank test, f-test was used as indicated in figure legends. P values of < 0.05 were considered significant. *P < 0.05, **P < 0.01, ***P < 0.001.

## Results

### Establishment of primary renal lymphoma model

The Eμ-Myc;Cdkn2a^-/-^ cell line, also called MA-LN in this study, was a cell line widely used to establish lymphoma transplantation model, a kind of GDA models. M referred to Myc gene and A referred to Arf gene. MA-LN Lymphoma typically presented in lymph nodes (LNs) in recipient mice **(Figure 1A)** and the progression of lymphoma could be well monitored by touching the palpable mass arising in the axillary lymph nodes.

**FIGURE 1.**
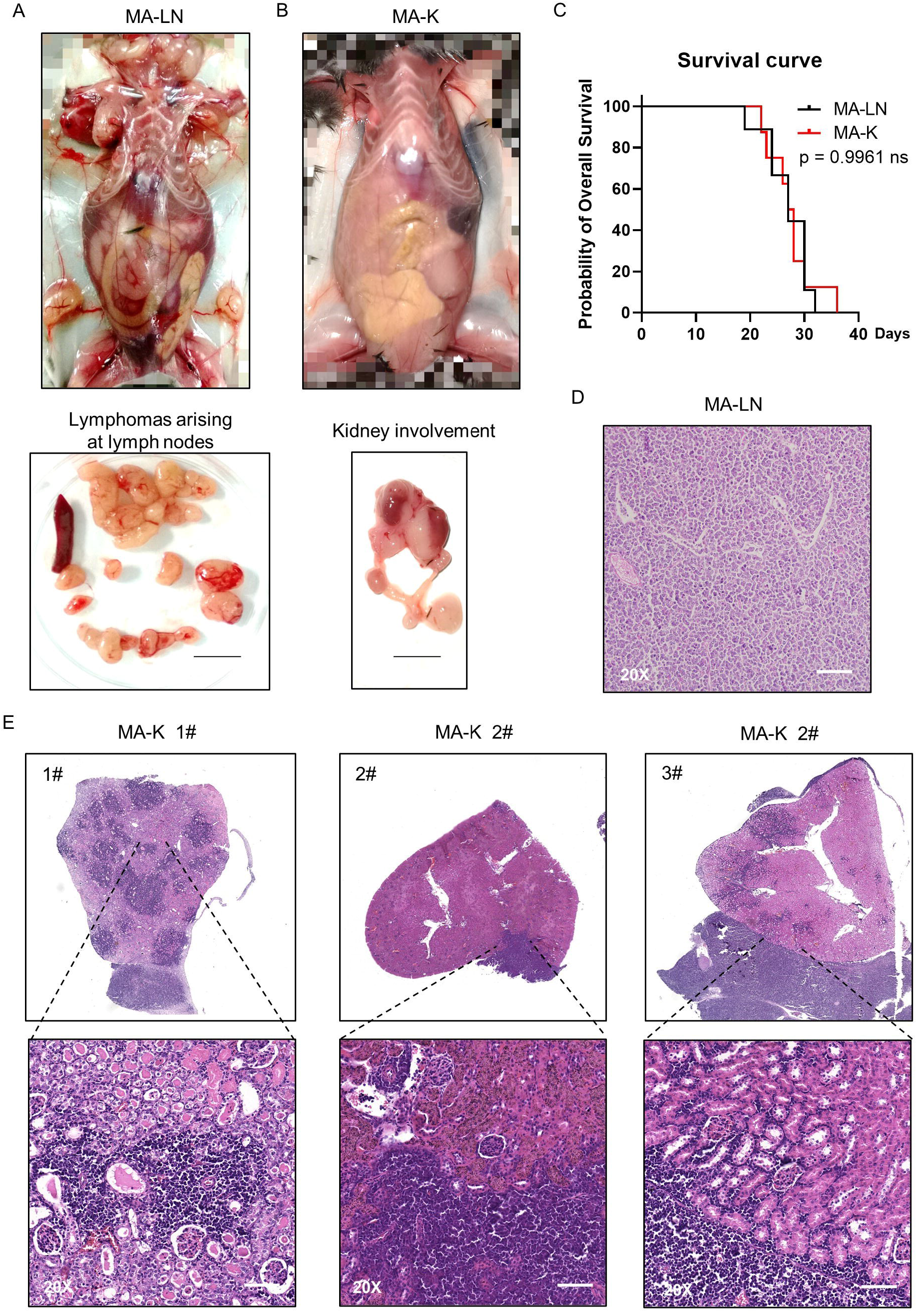
MA-K tends to disseminate in the kidney of recipient mice in lymphoma transplantation model. **(A)**. A representative picture showing the disseminated sites of MA-LN lymphoma in recipient mouse. The lymphomas were dissociated from mandibular lymph nodes, axillary lymph nodes, and inguinal lymph nodes of the recipient mice. Spleen was not enlarged. Bar, 1cm. **(B)**. A representative picture showing the unaffected lymph nodes and affected kidney and ureter in MA-K recipient mice. Bar, 1cm. **(C)**. Kaplan-Meier plots of MA-LN and MA-K recipient mice. n (MA-LN) = 9, n (MA-K) = 8. The equality of survival curves was tested using a log-rank test. **(D)**. A representative H&E staining section of MA-LN lymphoma. Bar in 20x image, 200 μm. **(E)**. Representative H&E staining sections of primary renal lymphoma arising in MA-K recipient mice. Bar in 20x image, 200 μm.

The MA-LN cell line was first introduced to established MA-LN GDA model in 2011. Two years ago, we began to notice that some obvious symptoms not seen before were appeared including hunch back, dull hair, movement retardation and abdominal bulge, instead of palpable mass at LNs. Anatomical results showed that lymphoma was mainly disseminated at the kidney and LNs were no longer involved **(Figure 1B)**, which was highly similar to primary renal lymphoma rarely in human. Despite the differences of disseminated sites, there was no difference in survival time of MA-LN and MA-K recipient mice **(Figure 1C)**. The result was understandable considering that the survival time was determined by the disease progression and therapeutic response.

Considering that we had changed the source of recipient mice, we suspected that the source difference of recipient mice probably contributed to kidney dissemination of lymphoma. Hence, we successively replaced recipient mice from 3 different sources. The results showed that lymphoma was still disseminated at the kidney. Hence, we ruled out the influence of recipient mice.

A study(22) had proven that cancer cell lines could evolve in culture, forming genetical and transcriptional heterogeneity and different drug response. Therefore, we proposed that the Eμ-Myc;Cdkn2a^-/-^ cell line had evolved into a novel and stable cell line, renamed as MA-K, indicating the tendency of MA-K to kidney dissemination in GDA model.

The histological analysis of MA-LN lymphoma showed that the lymphoma mass was mainly composed of lymphoma cells **(Figure 1D)**. For MA-K lymphoma, we analysis several affected kidneys and typical H&E staining sections were presented **(Figure 1E)**. We found that MA-K cells usually started to invade the kidney from the glomerulus and the later lymphoma could also invade the renal capsule.

### MA-K had specialized transcriptional patterns and abnormal expression of LymphGen

To identify the molecular characteristics of MA-K cells, we performed RNA-Seq analysis and 12809 genes were initially detected in MA-LN and MA-K. To obtain differentially expressed genes (DEGs) with biological significance, we removed genes that FPKM<1 and 8547 genes were left. Surprisingly, about 20% genes (1905 in 8547) had differentially expressed more than twice, indicating that MA-K was completely different from MA-N at transcriptional level. Given that cancer cell lines could transcriptional evolve in culture(22), we attributed the huge transcriptional difference to transcriptional selection and adaptation during the culture.

To identify the molecular patterns of MA-LN and MA-K, we proposed the specialized transcriptional pattern (STP) and specially expressed genes (SEGs), instead of routinely DEGs, to describe the molecular pattern of the individual sample. 4 reference genes, Gapdh, Actb, Hsp90ab1 and Myc, was used as the reference gene panel to the quality and comparability of FPKM data. 44 LymphGen genes were selected to perform STP analysis. Due to basic expression level (FPKM<1), Bcl2, Bcl6, Bcl10 and other LymphGen genes were not included in the panel of 44 LymphGen genes. Compared to MA-LN, 14 down-regulated SEGs and 3 up-regulated SEGs (log_2_(FC) <=1 and log_2_(FC) >=1) in MA-K were identified in 44 LymphGen genes **(Figure 2A)**. The observation indicated that gene inactivation by transcriptional inhibition was happened during the evolution of MA-K, which was in consistent with high frequency inactivation mutations in human B-NHL.

**FIGURE 2.**
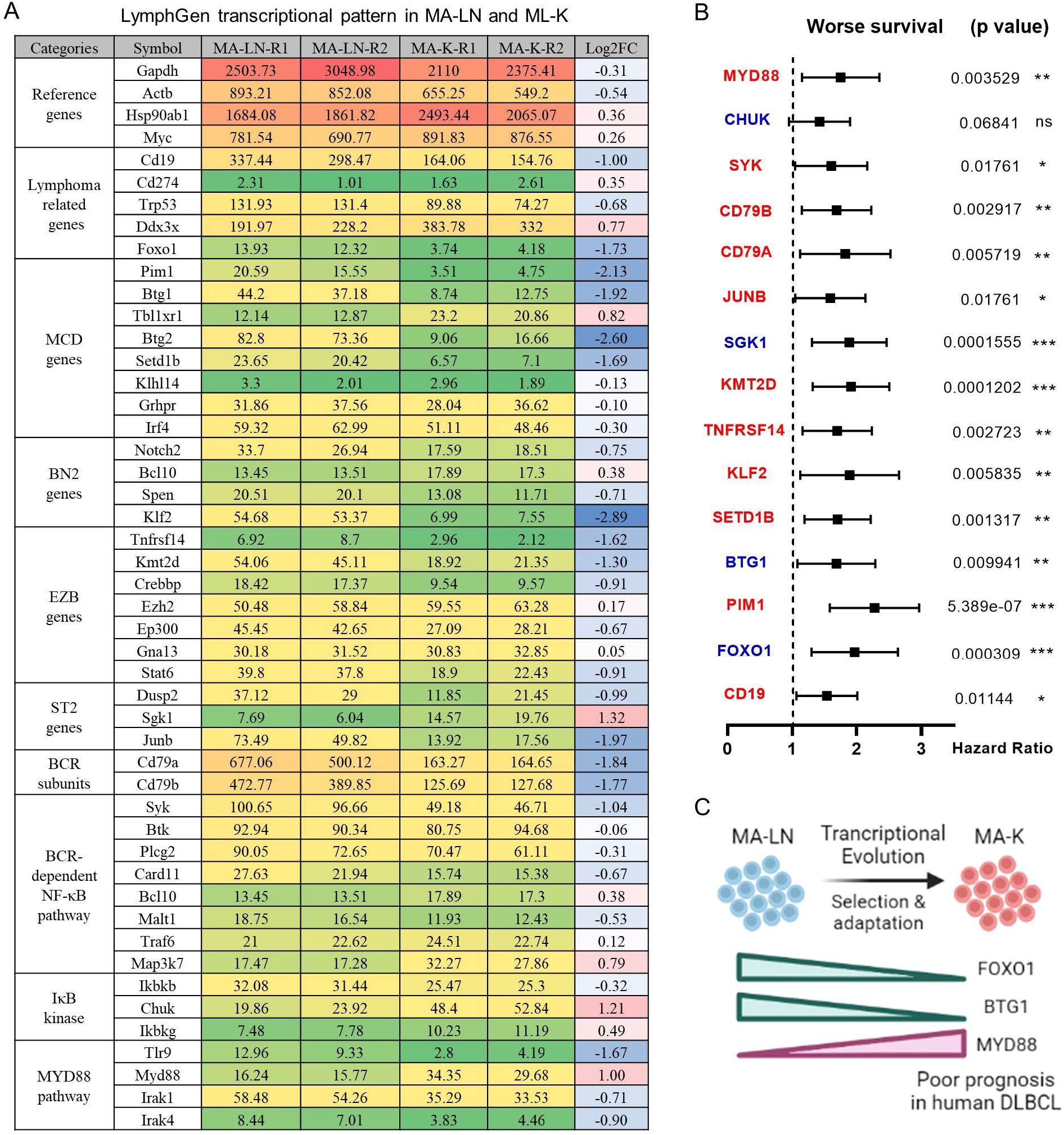
LymphGen signature in specialized transcriptional pattern of MA-K and clinical relevance. **(A)**. Expression level of LymphGen genes in MA-LN and MA-K. FPKM were used to evaluate the gene expression abundance in MA-LN and MA-K. The reference gene panel including Gapdh, Actb, Hsp90ab1 and Myc was presented to the quality and comparability of FPKM data. Log_2_(FC) was calculated by the average FPKM. FC, Fold Change. **(B)**. Forest plot of indicated genes in 2 risk groups. p-Value of the log-rank test were shown. The hazard ratio (HR), confidence interval, p-Value in forest plot were obtained from the SurvExpress program. **(C)**. Diagram for the formation of FOXO1-BTG1-MYD88 pattern during the evolution of MA-K. Created with BioRender.com.

To test whether gene expression level of SEGs in MA-K could predict clinical outcome, we performed survival analysis using a human DLBCL dataset (Lenz Staudt Lymphoma GSE10846, n = 420). We assumed that the STP of MA-K was associated with poor prognosis. The SurvExpress program was used to validate if the gene expression status could predict prognosis, in which the expression data of a single gene or multiple genes was calculated to the risk score, also called the prognostic index (PI)(20). The human DLBLC dataset (Lenz Staudt Lymphoma GSE10846, n = 420) was chosen, which contained detailed clinical information and had been adopted by many studies.

The results showed that most genes were associated with clinical relevance, while most of gene expression level in risk groups was contrary to expectations **(Figure 2B)**. Only FOXO1, BTG1, MYD88 were in line with expectations **(Figure 2C)**, indicating that abnormal transcriptional pattern for specialized lymphoma was very complicated due to the heterogeneity of lymphoma. Although the significance of transcriptional evolution of MA-K was not fully understood, most LymphGen genes were indeed significantly altered at transcriptional level.

### Translation pathway is altered in MA-K with clinical relevance

To filter SEGs with biological significance, we analyzed the expression data as follows **(Figure 3A)**. Inactive genes or basic expressed genes (defined as FPKM<1) in both MA-LN and MA-K had been excluded in the 8547 genes. For active genes (FPKM>=1) in both, genes with log_2_(FC) <=2 and log_2_(FC) >=2 were filtered as SEGs. In the case of inactive genes in one, active genes (FPKM>=10) in another were filtered as SEGs. All SEGs were listed in **Data Sheet 1**. A total of 360 SEGs were identified, specifically, 100 up-regulated SEGs and 260 down-regulated SEGs in MA-K.

**FIGURE 3.**
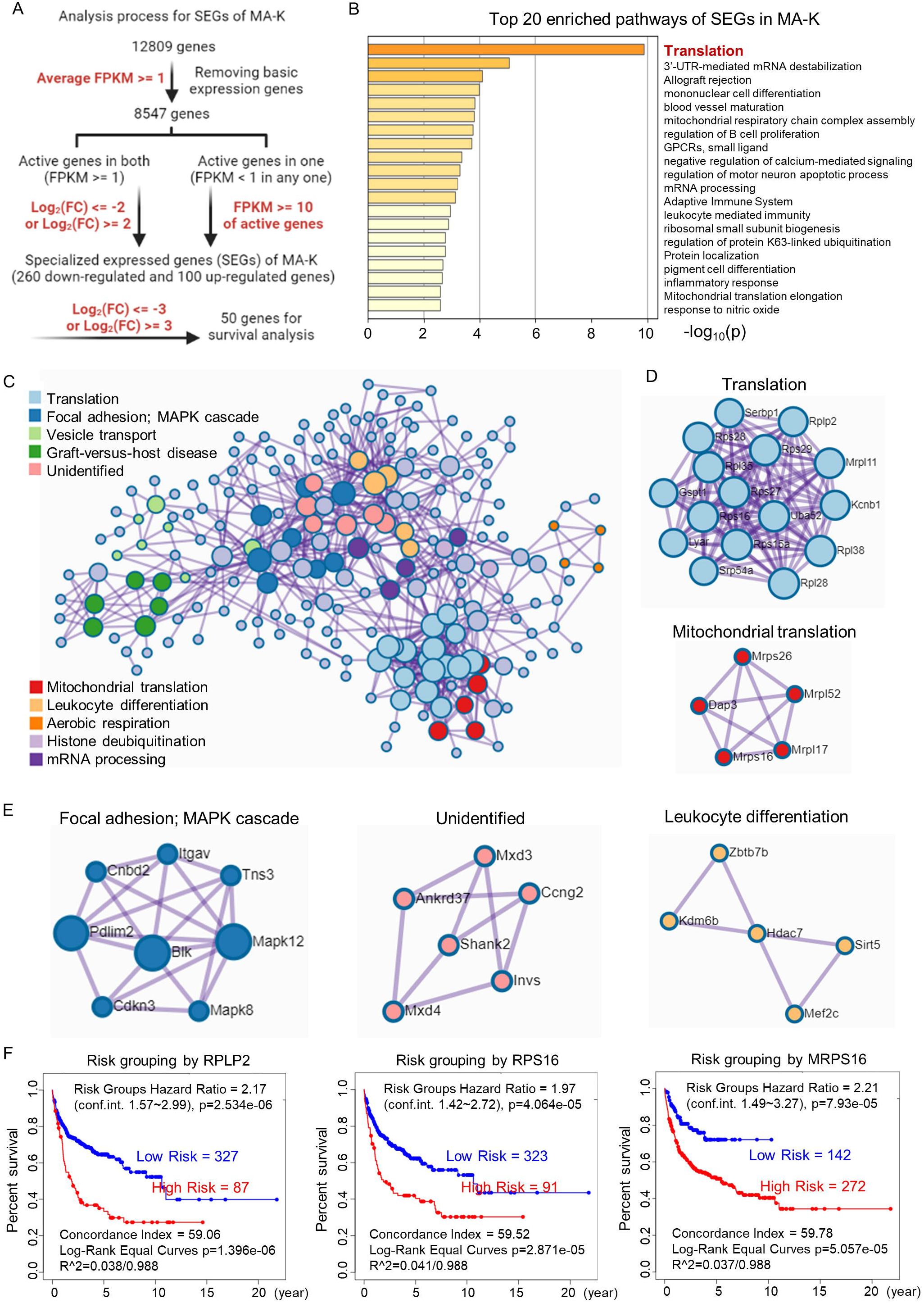
Translation related proteins, altered in MA-K, are associated with prognosis of human DLBCL. **(A)**. The analysis process of SEGs in MA-K. Filter parameters were highlighted in red. Inactive genes or basic expressed genes were defined as FPKM<1 and excluded in SEGs analysis. Active genes were defined as FPKM>=1. Created with BioRender.com. **(B)**. The top 20 enriched pathways of SEGs in MA-K. In total, 360 SEGs were selected as described in method and applied to pathway enrichment analysis. p-values are calculated based on the cumulative hypergeometric distribution. **(C)**. The protein-protein interaction (PPI) networks of SEGs. **(D)**. Genes in the PPI network including translation and mitochondrial translation. **(E)**. Genes in the PPI network including Focal adhesion-MAPK, Unidentified, Leukocyte differentiation. **(F)**. Kaplan-Meier plots of RPLP2, RPS16 and MRPS16 in human DLBCL. Red, high-risk group. Blue, low-risk group. Risk groups were generated based on the prognostic index (PI) for each gene and the optimization algorithm was applied in risk grouping. The number of each risk group was indicated in the plots. The equality of survival curves was tested using a log-rank test. A human DLBCL dataset (Lenz Staudt Lymphoma GSE10846, n = 420) was chosen to survival analysis.

The pathway enrichment analysis revealed that translation pathway was the most affected pathway in SEGs **(Figure 3B)**. In addition, the protein-protein interaction (PPI) network analysis discovered two core PPI networks in SEGs **(Figure 3C)**. A core PPI network involved proteins in the translation machine, including the mitochondrial translation machine **(Figure 3D)**. Another core PPI network involved proteins in Focal adhesion-MAPK, Unidentified, Leukocyte Differentiation **(Figure 3E)**. Together, the results suggested that the alteration of translation pathway and others presented the molecular features of MA-K cells.

Considering that many ribosomal proteins were abnormal regulated in tumors, we further analyzed the clinical relevance of the expression level of ribosomal proteins in human DLBCL. Survival analysis revealed that high expression level of RPLP2, RPS16 and MRPS16, up-regulated in MA-K, were significantly correlated with the poor prognosis of human DLBCL **(Figure 3F)**.

Together, the transcriptional profiles discovered that translation pathway and others were largely altered in MA-K. Survival analysis highlighted the clinical relevance of abnormal activation of ribosomal proteins in human DLBCL, indicating that drugs on targeting translation machine should be evaluated in MA-K GDA model and clinical trials. Whether abnormal activation of ribosomal proteins contributed to extranodal dissemination of lymphoma and other aggressive phenotypes should be further explored.

### Identification of 5-gene signature to predict prognosis of human DLBCL

Next, we investigated the clinical relevance of MA-K by evaluating the ability of SEGs in MA-K to predict prognosis of human DLBCL. Due to too many SEGs in MA-K, we only chosen 50 SEGs filtered with Log_2_(FC)<-3 or Log_2_(FC)>3 for survival analysis **(Figure 4A)**. Finally, 5 genes, down-regulated in MA-K to MA-L, SSPN, CD52, VPREB3, CCDC82, ANKRD37, were significantly correlated with poor prognosis of human DLBCL **(Figure 4B)**.

**FIGURE 4.**
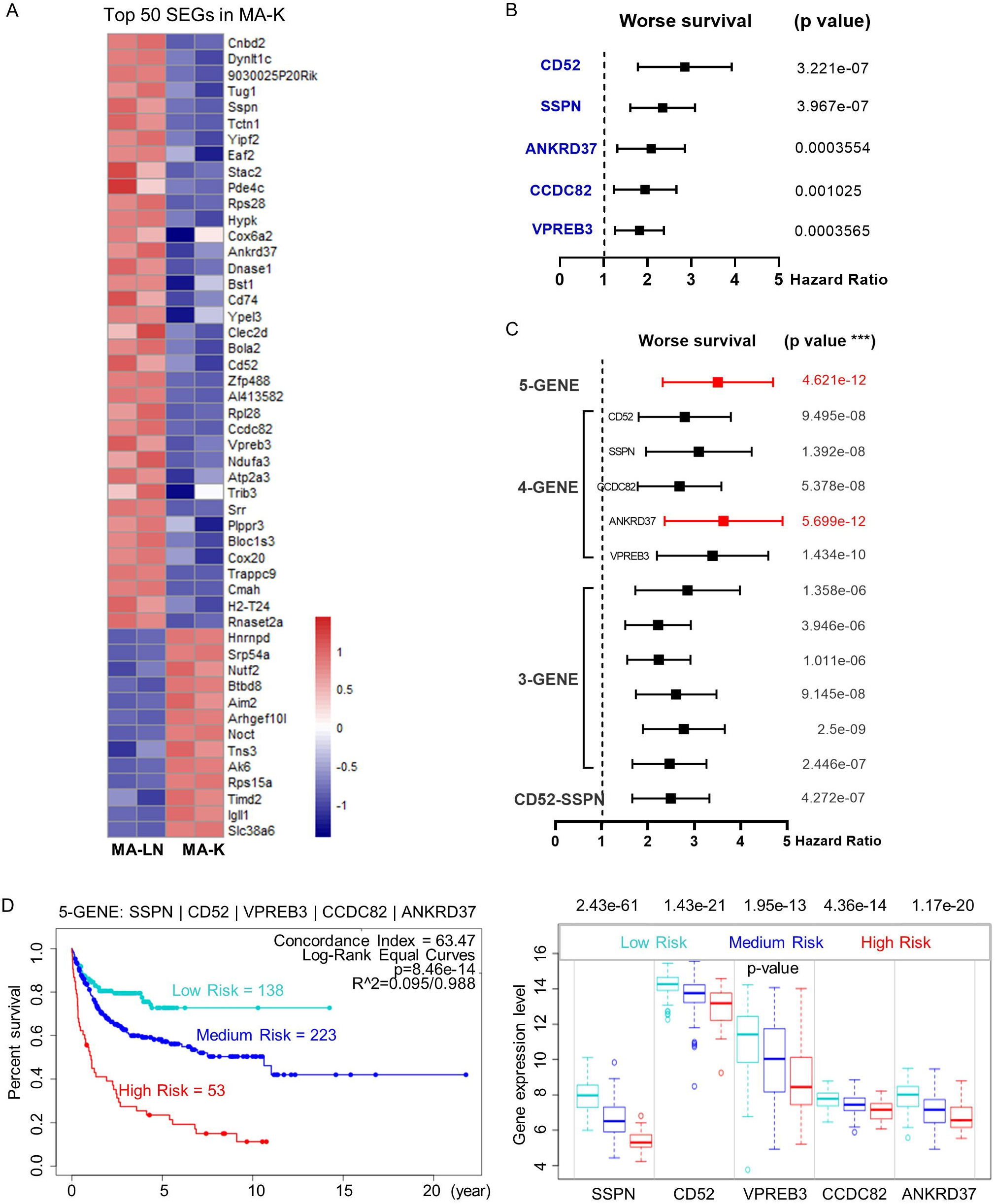
The 5-gene signature, altered in MA-K, predicts the prognosis of human DLBCL. **(A)**. Heatmap of SEGs in MA-K. 50 SEGs in MA-K (log_2_(FC) <=3 and log_2_(FC) >=3) were analyzed. FPKM of selected SEGs were used to generate the heatmap in R studio. **(B)**. Forest plot of SSPN, CD52, VPREB3, CCDC82, ANKRD37 in 2 risk groups. p-Value of the log-rank test were shown. The hazard ratio (HR), confidence interval, p-Value in forest plot were obtained from the SurvExpress program. **(C)**. Forest plot of different combinations of SSPN, CD52, VPREB3, CCDC82, ANKRD37 in 2 risk groups. p-Value of the log-rank test were shown. The hazard ratio (HR), confidence interval, p-Value in forest plot were obtained from the SurvExpress program. **(D)**. Kaplan-Meier plots of 5-gene signature and box plot of gene expression by 3 risk groups. Red, high-risk group. Cyan, medium-risk group. Blue, low-risk group. Risk groups were generated based on the prognostic index (PI) for each gene set and the optimization algorithm was applied in risk grouping. The number of each risk group was indicated in the plots. The p-valve of gene expression by 3 risk groups in box plot was obtained from an f-test. A human DLBCL dataset (Lenz Staudt Lymphoma GSE10846, n = 420) was chosen to survival analysis.

To investigate whether the 5 genes had a synergistic effect on the predicted clinical outcome, we compared the hazard ratio (HR) and p-Value of 3-gene and 4-gene combinations in 2 risk groups. Notably, the 5-gene signature showed improved prognostic prediction (HR=2.39, 95% confidence interval:3.37 to 4.76, p=4.621e-12) and a 4-gene signature that removed ANKRD37 was similar with the 5-gene signature, even better on HR. The results indicated that SSPN, CD52, VPREB3, CCDC82 were independent prognostic biomarkers and the 4-gene or 5-gene signature could be developed as a promising prognostic biomarker panel for human DLBCL **(Figure 4C)**.

To further evaluate the usefulness of the 5-gene signature, we tested its performance in 3 risk groups. The survival curves of high-, medium-, low-risk groups were well stratified by 5-gene signature (Log-Rank Equal Curves p=8.46e-14) and the individual genes were also significantly differential expressed in each risk groups **(Figure 4D)**. The p-Value of the 4-gene signature was a little worse than the 5-gene signature (Log-Rank Equal Curves p=1.465e-13).

We also noticed that the expression level of SSPN was the most different in 3 risk groups (p=2.43e-61). Given that SSPN was a membrane protein, it suggested that SSPN, detectable by immune-based assay, was a promising prognostic biomarker for human DLBCL. These results not only identified the 5-gene signature as a potential prognostic biomarker of human DLBCL, but also highlighted the clinical relevance between MA-K cell model and human aggressive DLBCL.

## Discussion

This study reported a novel murine cell line, named MA-K. Compared to parental Eμ-Myc;Cdkn2a^-/-^ cell line, called MA-LN in this study, MA-K cells tended to invade the kidney in lymphoma transplantation model, resembling primary renal lymphoma rarely in human. Kidney involvement was a kind of extranodal lymphoma and was associated with poor prognosis in human aggressive B-cell lymphoma. Although lymphoma arising in the kidney was rare(10), the remarkable behavior of MA-K that lymphoma did not present in lymph nodes in recipient mice was a notable feature for most kinds of human primary extranodal lymphoma. Therefore, MA-K could be developed as an important research model for primary extranodal lymphoma. MA-K cell line originated from MA-LN cell line and was identified as its kidney dissemination in recipient mice. Although we did not know how the MA-K cell line was formed, we could confirm that MA-K was largely different from the parent MA-LN at transcriptional level. Given that cancer cell lines could undergone the genetic and non-genetic evolution in culture, we attributed the formation of MA-K to transcriptional selection and adaptation (TSA)(22,23). We proposed that both the genetic mechanism, such as abnormal B-cell differentiation, and the non-genetic mechanism, such as cell plasticity at transcriptional selection and adaptation, were involved in the evolution of MA-K. The causes for the formation of MA-K needed to be further analyzed and the experimental verification.

To establish the relevance of MA-K and human aggressive B-cell lymphoma, we analyzed specialized transcriptional pattern in MA-K by 44 LymphGen genes. FOXO1 was frequently mutated in EZB-MYC+ subtype(2). BTG1 was frequently mutated in MCD subtype. Oncogenically active mutations in MYD88 were observed in many extranodal lymphoma of DLBCL (24–29) and classified into MCD subtype(2). FOXO1, BTG1, MYD88 were in line with expectations, indicating that MA-K shared the molecular pattern of MCD and EZB-MYC+ subtypes.

To confirm the clinical relevance of MA-K, we re-examined the SEGs in MA-K from two perspectives. In terms of signal pathway enrichment, we found that translation related ribosomal proteins were enriched in MA-K, and the high expression of these genes was also correlated to the poor prognosis of human DLBCL. Emerging evidence suggested that dysregulation of onco-ribosomes could facilitate the oncogenic translation program and increase the risk of developing malignancy, including tumor behavior, therapeutic response and clinical outcome(30–32). RPLP2, RPS16 and MRPS16, discovered in this study, had been reported to play oncogenic roles in various tumors(33–35). Hence, the gain of onco-ribosomes in MA-K suggested pharmaceutical inhibition of translation could be a potential therapeutic vulnerability of aggressive B-cell lymphoma. In terms of top SEGs, we chosen the top 50 SEGs for evaluation of prognostic biomarkers. The five genes, down-regulated in MA-K, were correlated with poor prognosis of human DLBCL, suggesting that these genes played negative regulation in aggressive progression of human DLBCL. Due to the limitation of gene expression datasets with clinical information, we only evaluated the clinical relevance between MA-K and human DLBCL.

Meanwhile, CD52 and SSPN, as membrane proteins, probably directly participated in the interaction between lymphoma cells and tumor microenvironment and finally determined lymphoma dissemination. In terms of molecular classification and molecular diagnosis, we proposed that CD52 and SSPN were ideal prognostic biomarkers to predicting the clinical outcomes. As a specific antigen in all blast cells, CD52 had been developed as a promising therapeutic target by mono-antibody (Alemtuzumab) in many clinical trials(36–39). However, CD52 was down-regulated in MA-K and the low expression of CD52 was correlated to poor prognosis of human DLBCL, suggesting that patients could not benefit from anti-CD52 immunotherapy and even worse. If CD52 was a negative regulator for aggressive B-cell lymphoma, targeting CD52 would directly accelerate malignant transformation of lymphoma, such as extranodal dissemination. Hence, the adoptation of anti-CD52 immunotherapy in clinical trials needed to be carefully reassessed.

In conclusion, the MA-K GDA model was a syngeneic lymphoma transplantation model, in which lymphoma arising in recipient mice usually disseminated at the kidney, highly resembling human primary renal lymphoma. SEGs in MA-K revealed that MA-K had strong clinical relevance with human aggressive DLBCL and onco-ribosomes and others, such as CD52 and SSPN, were identified as promising prognostic biomarkers in human DLBCL. Further studies on the MA-K cell line would provide more meaningful insights into the genetic and non-genetic mechanism of extranodal lymphoma. We hope that the MA-K GDA model could be widely used as preclinical model of aggressive B-cell lymphoma for efficacy evaluation of chemotherapy and immunotherapy.

## Supporting information

Data Sheet 1

## Data Availability Statement

All raw data generated or analyzed during this study are included in the article and in its supplementary material. RNA-Seq data of this study have been deposited in NCBI Sequence Read Archive (SRA) with accession codes PRJNA763338. The cell lines generated in this study can be acquired to the corresponding author with the Materials Transfer Agreement (MTA).

## Ethics statement

The animal study was reviewed and approved by the Animal Care and Use Committee of Laboratory Animal Research Center, Jiangsu University.

## Author Contributions

XXL conceptualized, designed the experiments and wrote the manuscript. HQ helped with designing the experiments and reviewing the manuscript. MYD and CXZ performed the mouse experiments. XXL analyzed the transcriptome data. LLL performed the prognostic analysis. All authors discussed the results and commented on the manuscript. XXL, HQ jointly supervised the study.

## References

1. Sehn LH, Salles G. Diffuse Large B-Cell Lymphoma. N Engl J Med (2021) 384:842–858. doi: 10/gh626w

2. Wright GW, Huang DW, Phelan JD, Coulibaly ZA, Roulland S, Young RM, Wang JQ, Schmitz R, Morin RD, Tang J, et al. A Probabilistic Classification Tool for Genetic Subtypes of Diffuse Large B Cell Lymphoma with Therapeutic Implications. Cancer Cell (2020) 37:551–568.e14. doi: 10/gjjd2m

3. Bociek RG. Adult Burkitt’s lymphoma. Clin Lymphoma (2005) 6: 11–20. doi: 10.3816/clm.2005.n.021

4. Blum KA, Lozanski G, Byrd JC. Adult Burkitt leukemia and lymphoma. Blood (2004) 104:3009–3020. doi: 10.1182/blood-2004-02-0405

5. Olszewski AJ, Kurt H, Evens AM. Defining and treating high-grade B-cell lymphoma, NOS. Blood (2022) 140:943–954. doi: 10.1182/blood.2020008374

6. Ok CY, Medeiros LJ. High-grade B-cell lymphoma: a term re-purposed in the revised WHO classification. Pathology (2020) 52:68–77. doi: 10.1016/j.pathol.2019.09.008

7. Schaff LR, Grommes C. Primary central nervous system lymphoma. Blood (2022) 140:971–979. doi: 10.1182/blood.2020008377

8. Willemze R, Jaffe ES, Burg G, Cerroni L, Berti E, Swerdlow SH, Ralfkiaer E, Chimenti S, Diaz-Perez JL, Duncan LM, et al. WHO-EORTC classification for cutaneous lymphomas. Blood (2005) 105:3768–3785. doi: 10.1182/blood-2004-09-3502

9. Cao X-X, Li J, Cai H, Zhang W, Duan M-H, Zhou D-B. Patients with primary breast and primary female genital tract diffuse large B cell lymphoma have a high frequency of MYD88 and CD79B mutations. Ann Hematol (2017) 96:1867–1871. doi: 10.1007/s00277-017-3094-7

10. Cheng X, Huang Z, Li D, Wang Y. Enormous primary renal diffuse large B-cell lymphoma: A case report and literature review. J Int Med Res (2019) 47:2728–2739. doi: 10.1177/0300060519842049

11. Alizadeh AA, Eisen MB, Davis RE, Ma C, Lossos IS, Rosenwald A, Boldrick JC, Sabet H, Tran T, Yu X, et al. Distinct types of diffuse large B-cell lymphoma identified by gene expression profiling. Nature (2000) 403:503–511. doi: 10/dqbqxp

12. Adams JM, Harris AW, Pinkert CA, Corcoran LM, Alexander WS, Cory S, Palmiter RD, Brinster RL. The c-myc oncogene driven by immunoglobulin enhancers induces lymphoid malignancy in transgenic mice. Nature (1985) 318:533–538. doi: 10/bv8xg4

13. Schmitt CA, McCurrach ME, Stanchina E de, Wallace-Brodeur RR, Lowe SW. INK4a/ARF mutations accelerate lymphomagenesis and promote chemoresistance by disabling p53. Genes Dev (1999) 13:2670–2677.

14. Eischen CM, Weber JD, Roussel MF, Sherr CJ, Cleveland JL. Disruption of the ARF–Mdm2–p53 tumor suppressor pathway in Myc-induced lymphomagenesis. Genes Dev (1999) 13:2658–2669.

15. Mori S, Rempel RE, Chang JT, Yao G, Lagoo AS, Potti A, Bild A, Nevins JR. Utilization of Pathway Signatures to Reveal Distinct Types of B Lymphoma in the Eμ-myc Model and Human Diffuse Large B-Cell Lymphoma. Cancer Res (2008) 68:8525–8534. doi: 10.1158/0008-5472.CAN-08-1329

16. Lefebure M, Tothill RW, Kruse E, Hawkins ED, Shortt J, Matthews GM, Gregory GP, Martin BP, Kelly MJ, Todorovski I, et al. Genomic characterisation of Eμ-Myc mouse lymphomas identifies Bcor as a Myc co-operative tumour-suppressor gene. Nat Commun (2017) 8:14581. doi: 10/f9svg3

17. Day C-P, Merlino G, Van Dyke T. Preclinical Mouse Cancer Models: A Maze of Opportunities and Challenges. Cell (2015) 163:39–53. doi: 10/f7tqq9

18. Zhao B, Pritchard JR, Lauffenburger DA, Hemann MT. Addressing Genetic Tumor Heterogeneity through Computationally Predictive Combination Therapy. Cancer Discov (2014) 4:166–174. doi: 10/gmcv8r

19. Ferrao PT, Bukczynska EP, Johnstone RW, McArthur GA. Efficacy of CHK inhibitors as single agents in MYC-driven lymphoma cells. Oncogene (2012) 31:1661–1672. doi: 10/ft58n7

20. Aguirre-Gamboa R, Gomez-Rueda H, Martínez-Ledesma E, Martínez-Torteya A, Chacolla-Huaringa R, Rodriguez-Barrientos A, Tamez-Peña JG, Treviño V. SurvExpress: an online biomarker validation tool and database for cancer gene expression data using survival analysis. PLoS One (2013) 8:e74250. doi: 10/ggcwpm

21. Lenz G, Wright G, Dave SS, Xiao W, Powell J, Zhao H, Xu W, Tan B, Goldschmidt N, Iqbal J, et al. Stromal gene signatures in large-B-cell lymphomas. N Engl J Med (2008) 359:2313–2323. doi: 10.1056/NEJMoa0802885

22. Ben-David U, Siranosian B, Ha G, Tang H, Oren Y, Hinohara K, Strathdee CA, Dempster J, Lyons NJ, Burns R, et al. Genetic and transcriptional evolution alters cancer cell line drug response. Nature (2018) 560:325–330. doi: 10.1038/s41586-018-0409-3

23. Marine J-C, Dawson S-J, Dawson MA. Non-genetic mechanisms of therapeutic resistance in cancer. Nat Rev Cancer (2020) 20:743–756. doi: 10/ghd84p

24. Ngo VN, Young RM, Schmitz R, Jhavar S, Xiao W, Lim K-H, Kohlhammer H, Xu W, Yang Y, Zhao H, et al. Oncogenically active MYD88 mutations in human lymphoma. Nature (2011) 470:115–119. doi: 10.1038/nature09671

25. Kraan W, van Keimpema M, Horlings HM, Schilder-Tol EJM, Oud MECM, Noorduyn LA, Kluin PM, Kersten MJ, Spaargaren M, Pals ST. High prevalence of oncogenic MYD88 and CD79B mutations in primary testicular diffuse large B-cell lymphoma. Leukemia (2014) 28:719–720. doi: 10.1038/leu.2013.348

26. Nakamura T, Tateishi K, Niwa T, Matsushita Y, Tamura K, Kinoshita M, Tanaka K, Fukushima S, Takami H, Arita H, et al. Recurrent mutations of CD79B and MYD88 are the hallmark of primary central nervous system lymphomas. Neuropathol Appl Neurobiol (2016) 42:279–290. doi: 10.1111/nan.12259

27. Cao X-X, Li J, Cai H, Zhang W, Duan M-H, Zhou D-B. Patients with primary breast and primary female genital tract diffuse large B cell lymphoma have a high frequency of MYD88 and CD79B mutations. Ann Hematol (2017) 96:1867–1871. doi: 10.1007/s00277-017-3094-7

28. Hattori K, Sakata-Yanagimoto M, Okoshi Y, Goshima Y, Yanagimoto S, Nakamoto-Matsubara R, Sato T, Noguchi M, Takano S, Ishikawa E, et al. MYD88 (L265P) mutation is associated with an unfavourable outcome of primary central nervous system lymphoma. Br J Haematol (2017) 177:492–494. doi: 10.1111/bjh.14080

29. Schrader AMR, Jansen PM, Willemze R, Vermeer MH, Cleton-Jansen A-M, Somers SF, Veelken H, van Eijk R, Kraan W, Kersten MJ, et al. High prevalence of MYD88 and CD79B mutations in intravascular large B-cell lymphoma. Blood (2018) 131:2086–2089. doi: 10.1182/blood-2017-12-822817

30. Elhamamsy AR, Metge BJ, Alsheikh HA, Shevde LA, Samant RS. Ribosome Biogenesis: A Central Player in Cancer Metastasis and Therapeutic Resistance. Cancer Res (2022) 82:2344–2353. doi: 10.1158/0008-5472.CAN-21-4087

31. Pelletier J, Thomas G, Volarević S. Ribosome biogenesis in cancer: new players and therapeutic avenues. Nat Rev Cancer (2018) 18:51–63. doi: 10.1038/nrc.2017.104

32. Bursać S, Prodan Y, Pullen N, Bartek J, Volarević S. Dysregulated Ribosome Biogenesis Reveals Therapeutic Liabilities in Cancer. Trends Cancer (2021) 7:57–76. doi: 10.1016/j.trecan.2020.08.003

33. Liao Y, Shao Z, Liu Y, Xia X, Deng Y, Yu C, Sun W, Kong W, He X, Liu F, et al. USP1-dependent RPS16 protein stability drives growth and metastasis of human hepatocellular carcinoma cells. J Exp Clin Cancer Res (2021) 40:201. doi: 10.1186/s13046-021-02008-3

34. Wang Z, Li J, Long X, Jiao L, Zhou M, Wu K. MRPS16 facilitates tumor progression via the PI3K/AKT/Snail signaling axis. J Cancer (2020) 11:2032–2043. doi: 10.7150/jca.39671

35. Artero-Castro A, Castellvi J, García A, Hernández J, Ramón y Cajal S, Lleonart ME. Expression of the ribosomal proteins Rplp0, Rplp1, and Rplp2 in gynecologic tumors. Hum Pathol (2011) 42:194–203. doi: 10.1016/j.humpath.2010.04.020

36. Geisler CH, van T’ Veer MB, Jurlander J, Walewski J, Tjønnfjord G, Itälä Remes M, Kimby E, Kozak T, Polliack A, Wu KL, et al. Frontline low-dose alemtuzumab with fludarabine and cyclophosphamide prolongs progression-free survival in high-risk CLL. Blood (2014) 123:3255–3262. doi: 10.1182/blood-2014-01-547737

37. Faderl S, Thomas DA, O’Brien S, Garcia-Manero G, Kantarjian HM, Giles FJ, Koller C, Ferrajoli A, Verstovsek S, Pro B, et al. Experience with alemtuzumab plus rituximab in patients with relapsed and refractory lymphoid malignancies. Blood (2003) 101:3413–3415. doi: 10.1182/blood-2002-07-1952

38. Uppenkamp M, Engert A, Diehl V, Bunjes D, Huhn D, Brittinger G. Monoclonal antibody therapy with CAMPATH-1H in patients with relapsed high- and low-grade non-Hodgkin’s lymphomas: a multicenter phase I/II study. Ann Hematol (2002) 81:26–32. doi: 10.1007/s00277-001-0394-7

39. Lundin J, Osterborg A, Brittinger G, Crowther D, Dombret H, Engert A, Epenetos A, Gisselbrecht C, Huhn D, Jaeger U, et al. CAMPATH-1H monoclonal antibody in therapy for previously treated low-grade non-Hodgkin’s lymphomas: a phase II multicenter study. European Study Group of CAMPATH-1H Treatment in Low-Grade Non-Hodgkin’s Lymphoma. J Clin Oncol (1998) 16:3257–3263. doi: 10.1200/JCO.1998.16.10.3257

